# Arabidopsis Mediator subunit 17 connects transcription with DNA repair after UV-B exposure

**DOI:** 10.1101/2021.08.02.454780

**Authors:** Marisol Giustozzi, Santiago Nicolás Freytes, Aime Jaskolowski, Micaela Lichy, Julieta Mateos, Maria Lorena Falcone Ferreyra, Germán L. Rosano, Pablo Cerdán, Paula Casati

## Abstract

Mediator 17 (MED17) is a subunit of the Mediator complex that regulates transcription initiation in eukaryotic organisms. In yeast and humans, MED17 also participates in DNA repair, physically interacting with proteins of the Nucleotide Excision DNA Repair system. We here analyzed the role of MED17 in Arabidopsis plants exposed to UV-B radiation, which role has not been previously described. Comparison of *med17* mutant transcriptome to that of WT plants showed that almost one third of transcripts with altered expression in *med17* plants are also changed by UV-B exposure in WT plants. To validate the role of MED17 in UV-B irradiated plants, plant responses to UV-B were analyzed, including flowering time, DNA damage accumulation and programmed cell death in the meristematic cells of the root tips. Our results show that *med17* and *OE MED17* plants have altered responses to UV-B; and that MED17 participates in various aspects of the DNA damage response (DDR). Increased sensitivity to DDR after UV-B in *med17* plants can be due to altered regulation of UV-B responsive transcripts; but additionally MED17 physically interacts with DNA repair proteins, suggesting a direct role of this Mediator subunit during repair. Finally, we here also show that MED17 is necessary to regulate the DDR activated by ATR, and that PDCD5 overexpression reverts the deficiencies in DDR shown in *med17* mutants. Together, the data presented demonstrates that MED17 is an important regulator of the DDR after UV-B radiation in Arabidopsis plants.

**One sentence summary:** In Arabidopsis, MED17 regulates the DNA damage response after UV-B exposure transcriptionally modulating the expression of genes and possibly also physically interacting with DNA repair proteins.

## Introduction

Mediator is a multi-subunit protein complex that regulates transcription initiation. Mediator acts as a molecular bridge between transcription factors bound at enhancers and RNA polymerase II (RNA pol II), but also regulates chromatin architecture, recruits epigenetic marks and participates in RNA processing. Structurally, Mediator is composed by four different modules, known as the head, middle, tail, and cyclin-dependent kinase 8 (CDK8) modules (reviewed in Buendía-Monreal and Gillmor, 2016; Malik et al., 2017; Mao et al., 2019). While the head module is thought to initially interact with RNA pol II to start transcription, the middle module has an important structural function and also binds to RNA pol II after its initial interaction with the head. The tail function is to associate with gene-specific transcription factors, whereas the CDK8 module is dissociable in response to different stimuli.

Even though the modular structure of the Mediator complex is conserved in eukaryotes, the composition of its subunits varies among species, but also changes in response to environmental and tissue-specific inputs, suggesting that different Mediator structures may have diverse functions (Mao et al., 2019). For instance, in plants, studies have shown that some Mediator proteins regulate cell division, cell fate and development, while others have a role in hormone signaling or are involved in biotic and abiotic stress responses. In plants, the Mediator complex comprises about 34 subunits (Malik et al., 2017). Despite mutations in different Mediator subunits are not lethal, altered expression of particular MED subunits can lead to important changes in gene expression. Therefore, med mutants show different phenotypes, for example they have altered growth, development and stress responses (Yang et al., 2016; Dolan and Chapple, 2017). For example, Arabidopsis mutants in *MED5*, *MED16* and *MED23* have decreased accumulation of phenylpropanoid compounds (Stout et al., 2008; Dolan et al., 2017); *med16* mutants are deficient in cellulose biosynthesis and iron homeostasis (Sorek et al., 2015; Yang et al., 2014; Zhang et al., 2014), while in *med15* and *cdk8* plants, lipid biosynthesis is altered (Kim et al., 2006; Zhu et al., 2014; Kong and Chang, 2018). Interestingly, several *med* mutants have shown contrasting metabolome profiles, which may relate to their different molecular function (Davoine et al., 2017).

MED17 is a subunit of the head Mediator module, and in the yeast and human complexes, it is an important protein which interacts with several other Mediator subunits (Guglielmi et al., 2004; Cevher et al., 2014). In Arabidopsis, MED17 seems to be a key scaffold component of the whole complex (Maji et al., 2019). AtMED17 was demonstrated to participate in the production of small and long noncoding RNAs (Kim et al., 2011). MED17 is required for small RNA biogenesis recruiting Pol II to promoters of miRNA genes, and also for the repression of heterochromatic loci, activating Pol II-mediated production of long noncoding RNAs (Maji et al., 2019). These results suggest that MED17 may have a role not only in transcription but also in genome stability. In yeast, MED17 participates in DNA repair through a physical interaction with Xeroderma pigmentosum group G protein (XPG), an endonuclease that participates in Nucleotide Excision DNA Repair (NER; Eyboulet et al., 2013). Moreover, *med17* mutants show increased sensitivity to UV radiation. Thus, MED17 participates in DNA repair recruiting XPG to transcribed genes (Eyboulet et al., 2013). In human cells, MED17 was shown to interact with the DNA helicase Xeroderma pigmentosum group B protein (XPB), which is a subunit of the transcription factor II H (TFIIH) and it is essential for both transcription and NER (Kikuchi et al., 2015). MED17 colocalizes with the NER factors XPB and XPG after UV-C exposure *in vivo*, and they also physically interact in *in vitro* assays, suggesting that, similarly as it was described in yeasts, MED17 plays essential roles in the switch between transcription and DNA repair (Kikuchi et al., 2015). The major DNA lesions that occur after UV exposure are the formation of cyclobutane pyrimidine dimers (CPDs) and pyrimidine (6-4) pyrimidones (6-4 PPs), which occur on two adjacent pyrimidine bases. In plants, these photoproducts are mainly repaired through photoreactivation, which is the direct reversal of major lesions by different types of photolyases that absorb light and reverse the formation of CPDs or 6-4 PPs (Spampinato, 2017). In Arabidopsis, UVR2 is a CPD photolyase, while UVR3 catalyzes the photoreactivation of 6-4 PPs. However, other DNA repair systems that are independent of light absorption, such as the NER system, which removes damaged nucleotides together with surrounding nucleotides; the Base Excision Repair (BER) system, which removes damaged bases, and other repair systems like the Mismatch Repair (MMR) system that recognizes and corrects of DNA mismatches have been also shown to repair DNA damage after UV exposure (Spampinato, 2017; Lario et al., 2011).

In this manuscript, we investigated the role of MED17 in Arabidopsis plants exposed to UV-B radiation. Using *med17* mutants, we analyzed their transcriptome and compared it to that of WT plants. Our results demonstrate that transcripts that encode proteins that participate in UV-B responses and DNA repair genes showed altered expression in *med17* mutants. *med17* mutants also showed altered phenotypes after UV-B exposure, in particular during the DNA damage response. We here show that MED17 is necessary for the correct expression of genes after UV-B exposure, probably by interacting with other Mediator proteins and transcription factors, but it also binds proteins that participate in DNA repair in Arabidopsis, suggesting that MED17 may have a direct role during DNA repair. In addition, MED17 is required for DDR activation through Ataxia telangiectasia and Rad3 related (ATR); and overexpression of Programmed cell death 5 (PDCD5), a regulator of the DDR in humans and in Arabidopsis, overcomes the deficiency of responses to UV-B in *med17* mutants. Together, the data presented here demonstrates that MED17 is a key regulator of the DNA damage response after UV-B radiation in Arabidopsis plants.

## Results

### Transcriptome analysis of *med17* mutants

As MED17 is a subunit of the transcriptional co-regulator Mediator complex, we were first interested in studying the global role of MED17 on gene expression. Thus, we compared the transcriptome of *med17* mutants and WT seedlings grown under white light for 10 days. Three biological replicates were used for RNA extraction and RNA-seq analysis. We identified 6822 differentially expressed genes in *med17* mutants in comparison to WT plants (Figure 1, A; Supplemental Table S1), 3534 (51.8%) were downregulated, whereas 3288 (48.2%) were upregulated. MED17 responsive genes were enriched in GO terms related to red, far-red, blue and UV-B light responses, among others (Supplemental Table S2). These results suggested a possible role for MED17 in UV responses. As described in the Introduction, MED17 was reported to participate in DNA repair after UV exposure in yeast and humans. Therefore, we compared the transcriptome changes of *med17* mutants to those of Col-0 plants grown under white light in our experiments (Supplemental Figure S1) to those previously reported for Col-0 plants after UV-B exposure (Tavridou et al., 2020). As shown in Figure 1, A, out of the 5079 UV-B regulated transcripts in WT plants (Tavridou *et al*., 2020), 2184 showed altered expression in *med17* mutants, which was significant by a Fisher Exact Test (p=4.6e-53). Interestingly, 56% of this overlapping set of transcripts were up-regulated by UV-B in WT plants and down-regulated in *med17* mutants (Supplemental Figure S1, A), which was highly significant (Fisher Exact Test p<1e-242; Figure 1, D). The overlap between up-regulated genes in *med17* mutants and down-regulated in Col-0 plants by UV-B was also significant, although not to the same extent (p <0.005; Figure 1, D). On the other hand, the overlap between the other two possible comparisons (up by UV-B vs up by *med17*; down by UV-B vs down by *med17* was 26 % altogether, and not statistically significant (Figure 1, D). Furthermore, the Odds ratio, which represents the strength of the association or correlation of the overlap, was higher for genes that are increased by UV-B and decreased in *med17* than for all the other comparisons (Figure 1, D). These results show that the transcripts that increase in response to UV-B require MED17 for maximal expression in the WT, suggesting a role for MED17 in UV-B responses.

**Figure 1.**
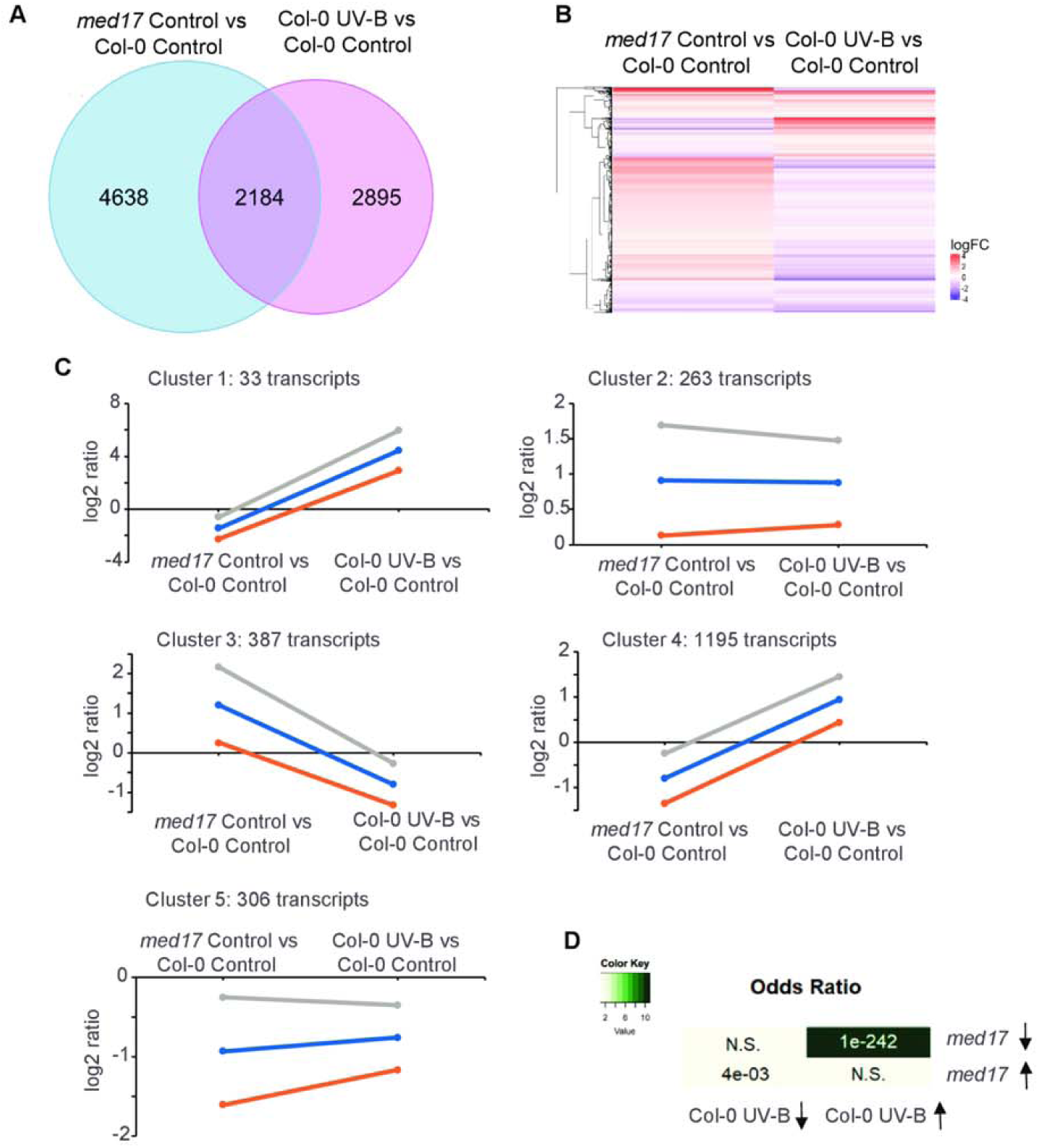
Analysis of global gene expression differences between *med17* compared to WT plants, and WT plants exposed to UV-B radiation. A, Venn diagram of comparisons between transcripts with altered expression in *med17* mutants and UV-B-responsive genes in Arabidopsis plants. Sets of genes were selected using the criteria described in Materials and Methods. B, Heatmap comparing transcripts changed in *med17* compared to WT plants; and WT plants after UV-B exposure. The color saturation reflects the magnitude of the log2 expression ratio for each transcript. C, Clusters of expression profiles. Each graph displays the mean pattern of expression of transcripts in the cluster in blue and the standard deviation of average expression (orange and grey lines). The number of transcripts in each cluster is at the top left corner of each graph. The *y*-axis represents log2 of gene-expression levels relative to those in WT Col-0 plants under control conditions without UV-B.

We then performed a cluster analysis of the 2184 overlapping genes shown in the Heatmap (Figure 1, B. Figure 1, C, and Supplemental Table S2). Genes were clustered in 5 groups, with clusters 1 and 4 including transcripts with decreased expression in *med17* compared to WT plants grown under white light conditions, and up-regulated by UV-B in Col-0 plants (Figure 1C). These two clusters mainly differ in the magnitude of the change observed; cluster 1 includes a lower number of transcripts, but showing stronger changes compared to WT plants, while cluster 4 includes more transcripts with smaller differences. Interestingly, when the clusters were analyzed by GO terms, cluster 1 included the categories UV-B responsive genes, flavonoid and secondary metabolites biosynthesis, and oxidoreduction reactions. This cluster contains highly upregulated genes in WT plants exposed to UV-B compared to plants under white light, but downregulated in *med17* mutants compared to Col-0, both grown in the absence of UV-B. Among genes included in cluster 1, we found *EARLY LIGHT-INDUCIBLE PROTEIN 2* (*ELIP2*), which was shown to be activated by UV-B through UVR8 (Brown, 2005), with a log2 fold change of 6.39 in WT under UV-B, and -3.29 in *med17* plants (Supplemental Table S2). *REPRESSOR OF UV-B PHOTOMORPHOGENESIS 2* (*RUP2*) was also found in this cluster. RUP2, along with RUP1, provides an UVR8 negative feedback regulation, balancing UV-B responses (Gruber, 2010). RUP2 showed a 3.20 log2 fold change in WT exposed to UV-B, while *med17* plants showed a log2 fold change of -1.45 (Supplemental Table S2). On the other hand, cluster 4 included GO terms related to light responses (light stimulus, high light, red, far-red, blue, photosynthesis), oxidative stress and response to reactive oxygen species, amongst others (Supplemental Table S2). Remarkably, in this cluster we found genes induced by UV-B like *DREB2A* (Ulm et al., 2004), and *RUP1*, mentioned above. This analysis further validates the requirement of MED17 for increased expression of genes that participate in UV-B responses.

### Photomorphogenic responses are altered in *med17* mutants

Because *med17* mutants had altered expression of some genes regulated by UV-B radiation, we studied UV-B responses in *med17* plants. First, we analyzed *med17* seedling lethality after UV-B exposure. Seedlings were grown on MS-agar plates under light conditions; a group of plants was irradiated with UV-B for 1h while a different group was kept in the dark for the same period. A third group of plants was grown under normal light conditions. Although *med17* seedlings germinated and grew similar to WT seedlings under white light illumination (100 μE m^−2^s^−1^); darkness affected the growth of some *med17* mutants. However, a UV-B treatment provoked lethality of most *med17* plants, while WT seedlings were very low affected by the same treatment (Figure 2, A).

**Figure 2.**
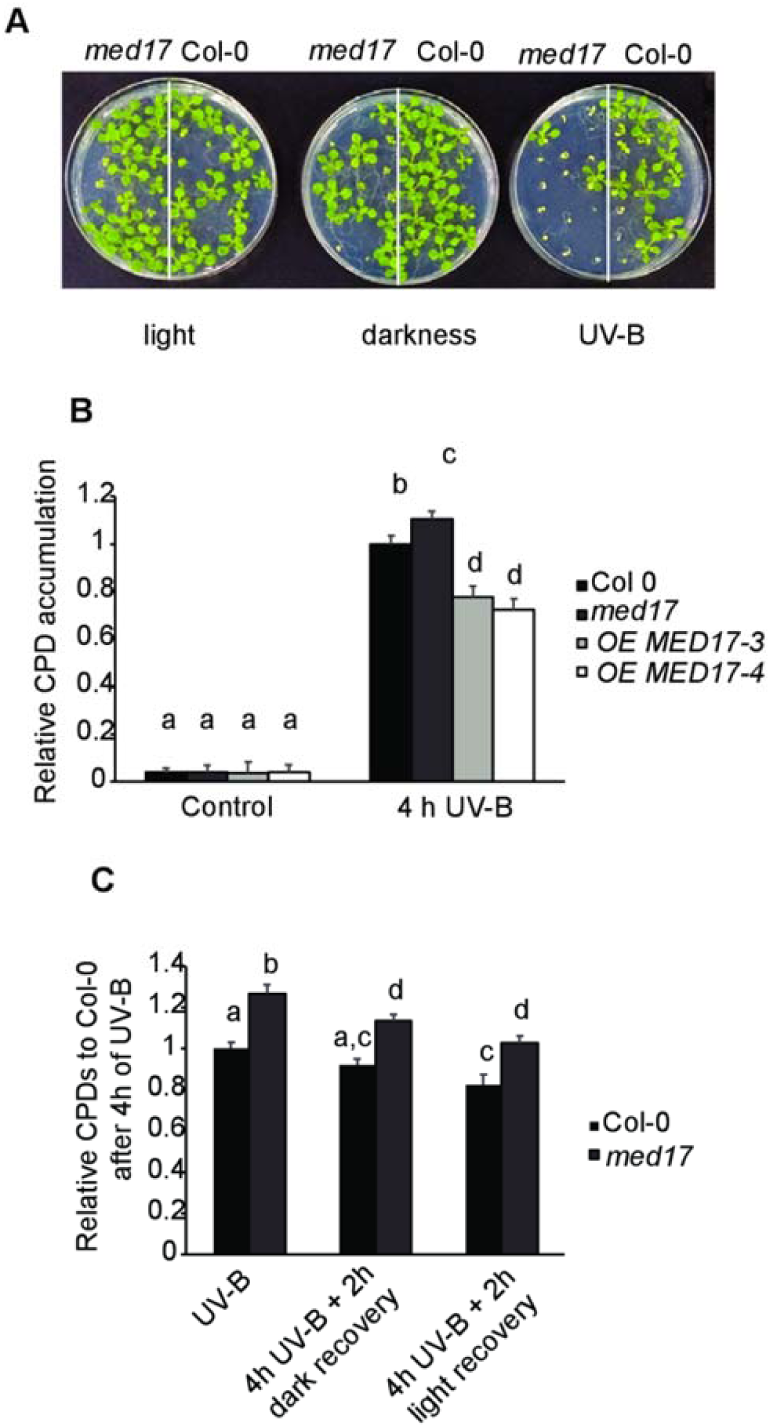
*med17* plants show higher UV-B sensitivity and DNA damage after UV-B than WT plants. A, Representative images of *med17* and WT Col-0 seedlings grown under light conditions and after 15 days were UV-B irradiated (UV-B) or kept under dark conditions (darkness) as described in Materials and methods. Alternatively, plants were grown under normal photoperiod (light conditions, and then kept under normal growth conditions after UV-B or kept under dark conditions. B and C, Relative CPD levels in the DNA of WT Col-0, *med17* and *OE MED17* plants grown under control conditions or immediately after a 4-h UV-B treatment under light conditions (B), or immediately after a 4-h UV-B treatment under dark conditions and 2 h after recovery in the dark or in the light to allow photorepair (C). Results represent averages ± S.E.M. of six independent biological replicates. Different letters indicate statistically significant differences applying ANOVA test (P <0.05).

Flowering time is delayed by UV-B radiation in Arabidopsis (Dotto et al., 2018; Arongaus et al., 2018). As previously reported in Kim et al. (2011) and similarly as it was also demonstrated for other Mediator subunit mutants such as *med25* and *med18* (Iñigo et al., 2012; Zheng et al., 2013), *med17* mutants flowered later than WT plants (Supplemental Figure S2, A-C). After growth under UV-B conditions, while WT plants showed a delay in flowering time, *med17* mutants showed a similar flowering time as that observed under control conditions. Thus, MED17 is a regulator of this developmental pathway and it may have a role in the regulation of flowering time under UV-B conditions. Together, the phenotypes analyzed show that *med17* mutants are sensitive to UV-B exposure.

### Plants with altered expression of *MED17* are deficient in the DNA damage response

We then focused on the role of MED17 in the DNA damage response after UV-B exposure. We first investigated if MED17 participates in DNA damage and repair using both *med17* mutants and transgenic plants that overexpress *MED17* under the control of the *35S* promoter (*OE MED17;* Supplemental Figure S3). While all plants showed very low and similar levels of CPDs under control conditions in the absence of UV-B, *med17* mutants accumulated higher DNA damage after a UV-B treatment than WT plants when the treatments were done under conditions that allowed photoreactivation (Figure 2, B). On the contrary, *OE MED17* plants had less CPDs after UV-B exposure under the same conditions. *med17* mutants also accumulated more CPDs than WT plants when the UV-B treatments were done in the dark to prevent DNA repair by photolyases, and 2h after the end of the UV-B treatment, either when recovery was done under light or dark conditions (Figure 2, C). Thus, *med17* mutants accumulate more CPDs after UVB exposure, and MED17 role during DNA damage and repair seems to be important not only during dark repair but also during photoreactivation.

We next analyzed whether plants with altered *MED17* expression showed differences in programmed cell dead (PCD) after UV-B exposure. When DNA damage occurs, DNA damage responses (DDRs) are triggered that converge to a PCD pathway, so as to avoid propagation of mutations in case the damage is not properly repaired (Furukawa et al., 2010). As shown in Figure 3, A, C, one day after a UV-B treatment, WT primary roots accumulated a higher number of dead cells after UV-B exposure than *med17* roots, and lower than *OE MED17* lines; while none of the analyzed lines showed any dead cells in untreated roots. However, 4 days after the treatment, although UV-B irradiated WT and *OE MED17* roots recovered and dead cells were almost undetectable; the number of dead cells in UV-B irradiated *med17* mutants was higher than in WT roots (Figure 3, B, C). Thus, MED17 is required for a proper activation of the PCD pathway after UV-B exposure.

**Figure 3.**
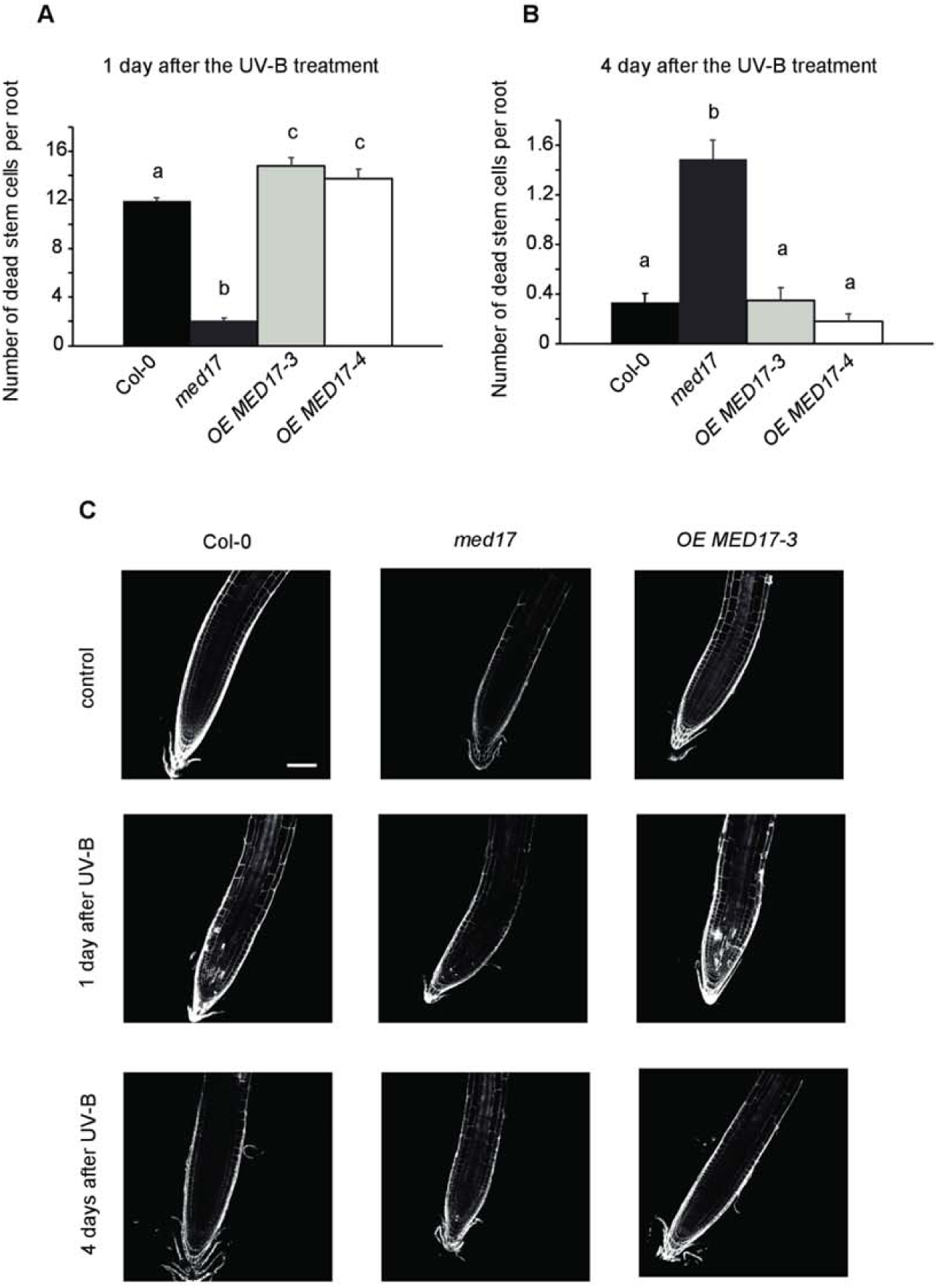
Programmed cell death in meristematic root cells in WT Col-0, *med17* and *OE MED17* plants after UV-B exposure. A and B, Number of stem cells that are dead after 1 day (A) or 4 days (B) of UV-B exposure in WT Col-0, *med17* and *OE MED17* roots. Results represent the average of at least 50 biological replicates ± S.E.M. Different letters indicate statistically significant differences applying analysis of variance test (*p* <0.05). C, Representative images of stem cells and adjacent daughter cells from WT Col-0, *med17* and *OE MED17* seedlings that were scored for intense PI staining to count dead stem cells per root 1 day and 4 days after a UV-B treatment or under control conditions. Scale bar represents 100 µm.

Another consequence of the DDR in plants is the inhibition of cell proliferation (Culligan et al., 2006). Thus, we analyzed the effect of UV-B on cell proliferation in the primary root meristems of plants with altered *MED17* expression. *med17* mutants had a smaller meristematic zone in the primary root than WT plants, which was significantly decreased after a UV-B treatment (Supplemental Figure S4, A and 5). The decrease in the meristematic zone size was higher in *med17* than in WT plants, in contrast to *OE MED17* plants, which were less inhibited by UV-B (Supplemental Figure 4, A and B). The higher decrease in the meristem size of *med17* roots by UV-B was a consequence of a higher inhibition of cortex cell proliferation than that measured in WT roots; and the opposite was observed in *OE MED17* roots (Supplemental Figure 4, C and D), while the increase in cortex cell length determined in the meristems of all plants by UV-B was similar (Supplemental Figure 4, E and F). In this way, MED17 also regulates cell proliferation after UV-B exposure, altering root meristem size.

### *med17* and *OE MED17* plants show altered UV-B regulation of gene expression

Next, we investigated how the expression of genes that respond to this radiation and regulate its responses was affected in *med17* seedlings after UV-B exposure. First, we analyzed the expression of three transcripts encoding enzymes that participate in DNA repair by UV-B, *UVR2* and *UVR3,* which encode CPD and 6-4 photoproduct photolyases, respectively; and *UVR7* (or *ERCC1*), which encodes a DNA repair endonuclease of the NER system. *UVR3* belongs to cluster 4 in Figure 1, C, showing decreased expression in *med17* plants (Supplemental Table S2). While the expression of *UVR2* was similar in *med17* and WT seedlings after UV-B exposure; *UVR3* and *UVR7* showed significantly lower levels than WT plants under control and UV-B conditions (Figure 4, A-C). Thus, the higher accumulation of CPDs by UV-B in *med17* plants could be due to decreased expression of DNA repair enzymes. As flowering time was also affected in *med17* plants, we analyzed the expression of *FLC*, which encodes a master repressor of flowering time (Michaels and Amasino, 1999). *FLC* levels were significantly higher in *med17* both under control conditions and after UV-B, with similar levels under both conditions (Figure 4, D); hence, the delay in flowering time in *med17* under both conditions could be explained by increased levels of this protein. Transcript levels of the DNA damage response kinases ATR and ATM, and the transcription factor SOG1, a master regulator of the DDR (Furukawa et al., 2010), were also analyzed. Fig 4, E-G, shows that both *ATR* and *ATM* levels were higher in *med17* than in WT plants, both under control conditions and after UV-B exposure. However, *SOG1* expression was significantly decreased in *med17* under both conditions analyzed, suggesting that decreased levels of this transcription factor may affect the DNA responses after UV-B exposure. On the contrary, *MAPK6* levels in *med17* mutants were not different to those in WT plants (Figure 4, H), this kinase is also a regulator of some UV-B responses, so the phenotypes observed in the mutants are not due to changes in the expression of this enzyme.

**Figure 4.**
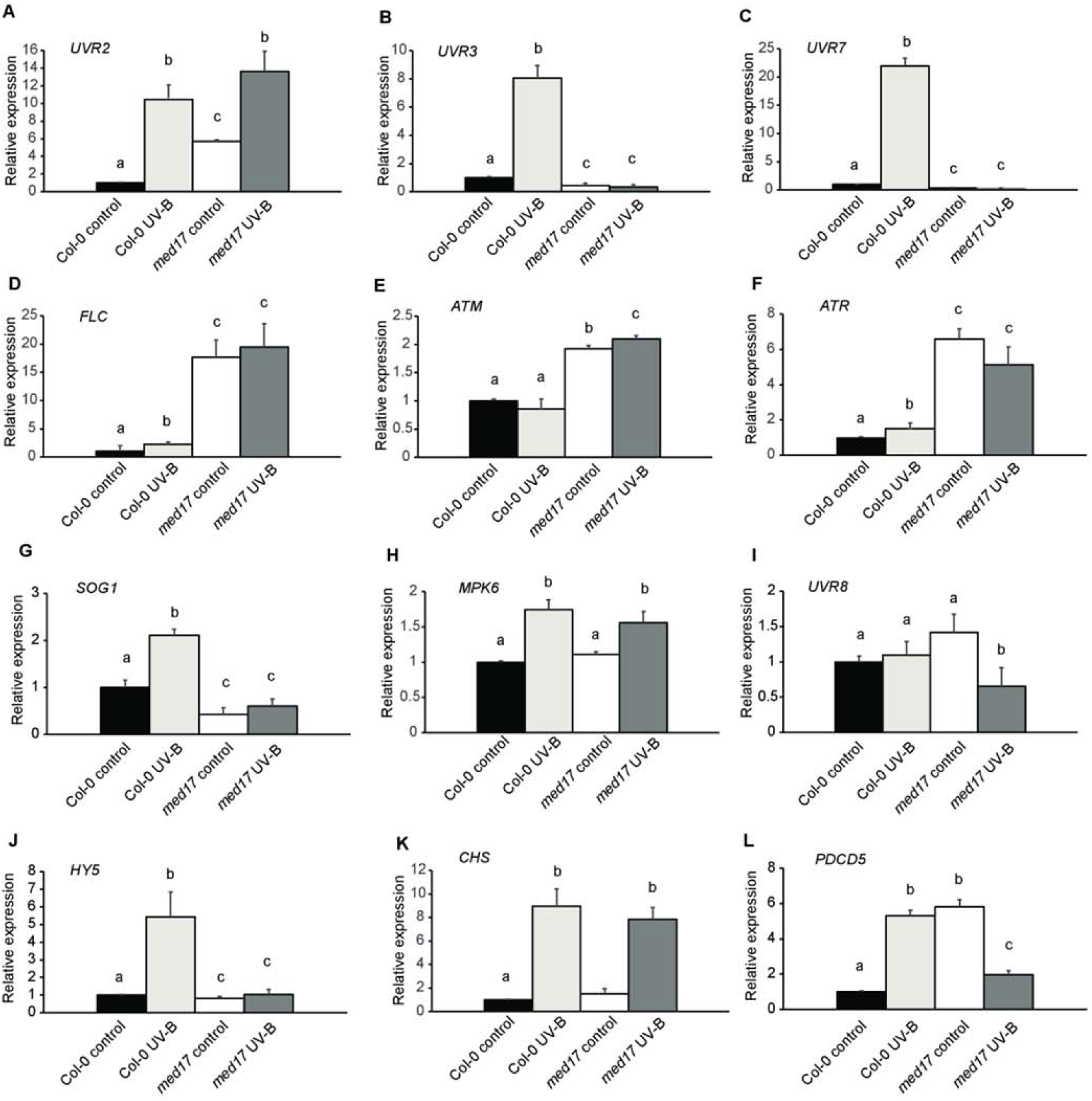
UV-B effect on expression of genes that participate in UV-B responses in WT Col-0 and *med17* seedlings. Relative expression levels of *UVR2* (A), *UVR3* (B), *UVR7* (C), *FLC* (D), *ATM* (E), *ATR* (F), *SOG1* (G), *MAPK6* (H), UVR8 (I), *HY5* (J), *CHS* (K) and *PDCD5* (L) analyzed by RT-qPCR in WT Col-0 and *med17* seedlings under control conditions or immediately after a 4 h-UV-B treatment (UV-B). Results represent the average ± SEM. Different letters indicate statistically significant differences applying an ANOVA test (P < 0.05). Data represent at least three biological replicate experiments. Each RT-qPCR was repeated at least three times on each biological replicate.

*UVR8*, which encodes the UV-B photoreceptor that regulates mostly photomorphogenic responses and which levels were not changed by UV-B in WT plants (Figure 4, I), was similarly expressed in *med17* and WT seedlings under control conditions, but showed a small but significant repression after UV-B exposure in the mutant. *HY5,* which encodes a transcription factor that regulates UV-B responses in the UVR8 pathway and showed decreased expression in *med17* plants in the RNA seq data (Supplemental Table S1), also showed decreased levels after UV-B exposure (Figure 5, J). Thus, under UV-B conditions, *med17* mutants have decreased expression of important regulators of the UV-B photomorphogenic pathway. Finally, *CHALCONE SYNTHASE* (*CHS*) transcript levels in *med17* were analyzed. *CHS* is a target of HY5 and encodes the first enzyme in the flavonoid pathway; these specialized metabolites provide UV-B protection in plants (Falcone Ferreyra et al., 2012). *CHS* expression was not altered in *med17* mutants, neither under control conditions nor after exposure (Figure 4, K). Interestingly, other transcripts that encode enzymes in the flavonoid pathway, such as *FLAVONOL SYNTHASE 1*, *CHALCONE ISOMERASE 1 and 3*, *FLAVONOID 3’-MONOOXYGENASE* (*CYP75B1*); and the transcription factor *MYB 111*, which regulates the expression of enzymes in the flavonoid pathway, belong to cluster 1 in the RNAseq data (Figure 1, B, Supplemental Table S1). These transcripts show decreased expression in *med17* mutants under control conditions. Despite this, flavonoid levels in *med17* and *OE MED17* plants were similar to those in WT plants (Supplemental Figure S2, D), correlating with the expression patterns of *CHS*. In this way, increased sensitivity to UV-B in *med17* plants is not due to changes in flavonoid accumulation.

**Figure 5.**
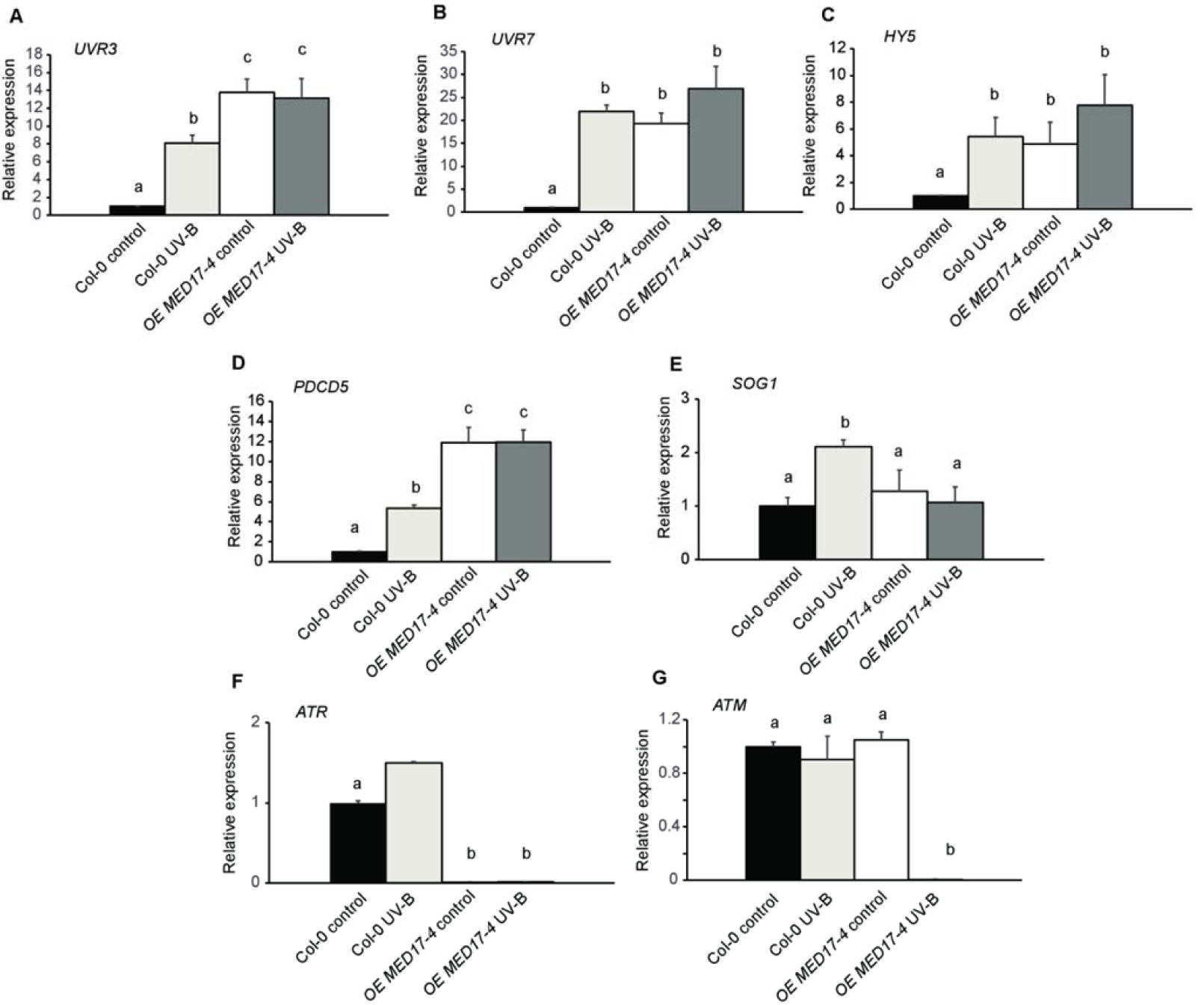
UV-B effect on expression of genes that participate in UV-B responses in WT Col-0 and *OE MED17* seedlings. Relative expression levels of *UVR3* (A), *UVR7* (B), *HY5* (C), *PDCD5* (D), *SOG1* (E), *ATR* (F) and *ATM* (G) analyzed by RT-qPCR in WT Col-0 and *OE MED17-4* seedlings under control conditions or immediately after a 4 h-UV-B treatment (UV-B). Results represent the average ± SEM. Different letters indicate statistically significant differences applying an ANOVA test (P < 0.05). Data represent at least three biological replicate experiments. Each RT-qPCR was repeated at least three times on each biological replicate.

On the other hand, when transcript levels of DNA repair enzymes with altered expression in *med17* mutants were analyzed in *OE MED17* plants, results showed that *UVR3* and *UVR7* were highly expressed in the transgenic plants, both under control conditions and after UV-B exposure (Figure 5, A, B), suggesting that lower CPD levels in *OE MED17* plants after UV-B exposure are due to high expression of these DNA repair enzymes. Interestingly, UV-B up-regulation of both genes is lost in the *OE MED17* plants; and this is also true for other UV-B responsive genes such as *HY5* and *SOG1* (Figure 5, C-E). Therefore, MED17 could mediate UV-B regulation of at least some UV-B marker genes, this may be through its regulation of *HY5* expression. On the contrary, both *ATR* and *ATM*, which were expressed at high levels in *med17* mutants, showed very low levels in *OE MED17* plants, in particular after UV-B exposure (Figure 5, F, G); thus, MED17 is a negative regulator of both DDR kinases.

Together, qRT-PCR results demonstrate that, for at least some genes, in particular *HY5*, *UVR3*, and *UVR7*, the upregulation after UV-B is lost in *med17* and constitutively increased in *OE MED17* seedlings; therefore, at least some of the phenotypes observed in UV-B irradiated plants could be due to altered expression these genes.

### MED17 interacts with nuclear proteins with roles during DNA repair

As described in the Introduction, in humans and yeasts, MED17, besides being a transcriptional regulator, also physically interacts with proteins that participate in DNA repair through the NER system. Therefore, to analyze if in Arabidopsis there is also a physical interaction of MED17 with DNA repair enzymes after UV-B exposure, Arabidopsis nuclei were obtained from UV-B irradiated *med17* mutants expressing *MED17* fused to *GFP* under the *35S* promoter. Expression of this fusion protein complemented the mutant phenotypes suggesting it is active *in vivo* (Supplemental Figure S3).

MED17 was co-immunoprecipitated from purified nuclei using anti-GFP antibodies, and the output was analyzed by LC-MS/MS (Smaczniak et al., 2012). Sixty-five nuclear proteins coimmunoprecipitated with MED17-GFP in at least 2 of 3 biological replicates from UV-B treated plants (Supplemental Table S3). In this group, there were other Mediator proteins, such as MED8, MED37 A/B, C and F; transcription initiation and splicing factors, chromatin associated proteins and other transcription factors (Supplemental Table S3). Interestingly, several proteins which were previously described to have a role during DNA repair were identified, such as the cohesin factor PDS5C (Pradillo et al., 2015), the histone chaperones NAP 1; 3 and NRP 1 and 2 (Casati and Gomez, 2021), two DEK domain-containing chromatin associated proteins (Waidmann et al., 2014), a sister chromatid cohesion 1 protein 4 (SYN4; da Costa-Nunes et al., 2004); a DNA repair ATPase-related protein and a replication factor C subunit 4 (Chen el al., 2018; Table 1, Supplemental Table S3). LC-MS data showed that MED17 is in a same complex with the transcription initiation factor TFIID subunit 9 and the transcription initiation factor IIE subunit alpha; therefore, MED17, together with these proteins, may be required for correct TCR repair as previously described in other species. Moreover, a cell division cycle 5-like protein (CDC5) was also found to immunoprecipitate with MED17 in plants exposed to UV-B, this protein has a role in cell cycle control and also in the response to DNA damage (Table 1; Lin et al., 2007). Thus, MED17 may not only directly participate in DNA repair, but it may also regulate the DNA damage response by interacting with other proteins acting downstream in the DDR pathway. This data suggests that, as demonstrated in yeasts and humans, AtMED17, besides having a role in transcription regulation by UV-B as shown in Figures 4 and 5, it could have a direct role interacting with nuclear proteins during DNA repair.

**Table 1.** Proteins with a putative DNA repair role enriched in the MED17-GFP IP experiments.

| Protein ID | Protein name | Accession number |
| --- | --- | --- |
| Q93ZX1 | Replication factor C subunit 4 (RFC4) | At1g21690 |
| Q94K07 | Nucleosome assembly protein 1;3 (NAP1;3) | At5g56950 |
| Q9CA59 | NAP1-related protein 1 (NRP1) | At1g74560 |
| Q8GUP3 | Precocious Dissociation of Sisters 5 (ATPDS5C) | At4g31880 |
| P92948 | Cell division cycle 5-like protein (CDC5) | At1g09770 |
| F4K4Y5 | DEK domain-containing chromatin associated protein | At5g55660 |
| Q9SUA1 | DEK domain-containing chromatin associated protein (DEK 3) | At4g26630 |
| Q8LC68 | NAP1-related protein 2 (NRP 2) | At1g18800 |
| Q8W1Y0 | Sister chromatid cohesion 1 protein 4 (SYN 4) | At5g16270 |
| Q9ZQ26 | DNA repair ATPase-related | At2g24420 |
| Q9SYH2 | Transcription initiation factor TFIID subunit 9 (TAF9) | At1g54140 |
| Q93ZW3 | Transcription initiation factor TFIIE subunit alpha | At1g03280 |

### *atr* mutant phenotypes are suppressed in the absence of MED17

To further analyze the role of MED17 in the DDR, we crossed *med17* mutants with either *atm* or *atr* plants to generate the corresponding double mutants. Ataxia telangiectasia mutated (ATM) is usually activated by double-strand breaks in the DNA; while ATR is mainly triggered by single-strand breaks or stalled replication forks such as CPDs and 6-4PPs that occur after UV-B exposure; and both independently regulate the DNA damage response in plants (Culligan et al., 2006). For the *med17 atm* crosses, only heterozygous mutants in either or both genes were obtained after screening 60 plants, but no double homozygous mutant plants were recovered, suggesting that *med17* deficiency generates lesions that require ATM to allow cell viability. On the contrary, *med17 atr* double homozygous mutants were obtained (Supplemental Figure S6). *med17 atr* plants were smaller than WT and *atr* plants when grown under standard growth conditions in the growth chamber, but they looked like *med17* mutants (Supplemental Figure S6, A). As *med17* single mutants, the double mutants had smaller leaves than WT and *atr* plants (Supplemental Figure S6, A). In addition, both the single *med17* and double *med17 atr* mutants displayed a significant reduction in the siliques length compared to WT and *atr* plants (Supplemental Figure S6, B and C). The number of seeds per silique in *med17* and *med17 atr* mutants was also decreased compared to WT and *atr* (Supplemental Figure S6, D). When the seeds in each silique were observed, *med17* and *med17 atr* mutants showed both fertilized seeds and aborted embryos, which correlates with the failure observed in seed production (Supplemental Figure S6, B, E and F). In contrast, both WT and *atr* mutants showed low and similar number of aborted seeds (Supplemental Figure S6, E and F). In order to analyze if embryos had dead cells, seeds were stained with propidium iodide and analyzed by confocal microscopy (Supplemental Figure S7). While WT and *atr* seeds showed a very low number of stained cells, both *med17* and *med17 atr* seeds showed higher staining. These results could explain the decreased fertility observed in *med17* and *med17 atr* mutants.

When plants were exposed to UV-B radiation, all mutants accumulated similar levels of CPDs and higher than those in WT plants (Figure 6, A). On the other hand, PCD was analyzed in the meristematic zone of the primary roots one day after a UV-B treatment and *med17 atr* mutants showed significantly fewer dead cells than *atr* and WT plants (Supplemental Figure S8). Moreover, 4 days after the treatment, *atr* mutants still had higher number of dead cells than the other lines under study, while *med17* and *med17 atr* mutants had a similar number of dead cells and higher than those in WT plants, which were almost recovered (Figure 6, B and C).

**Figure 6.**
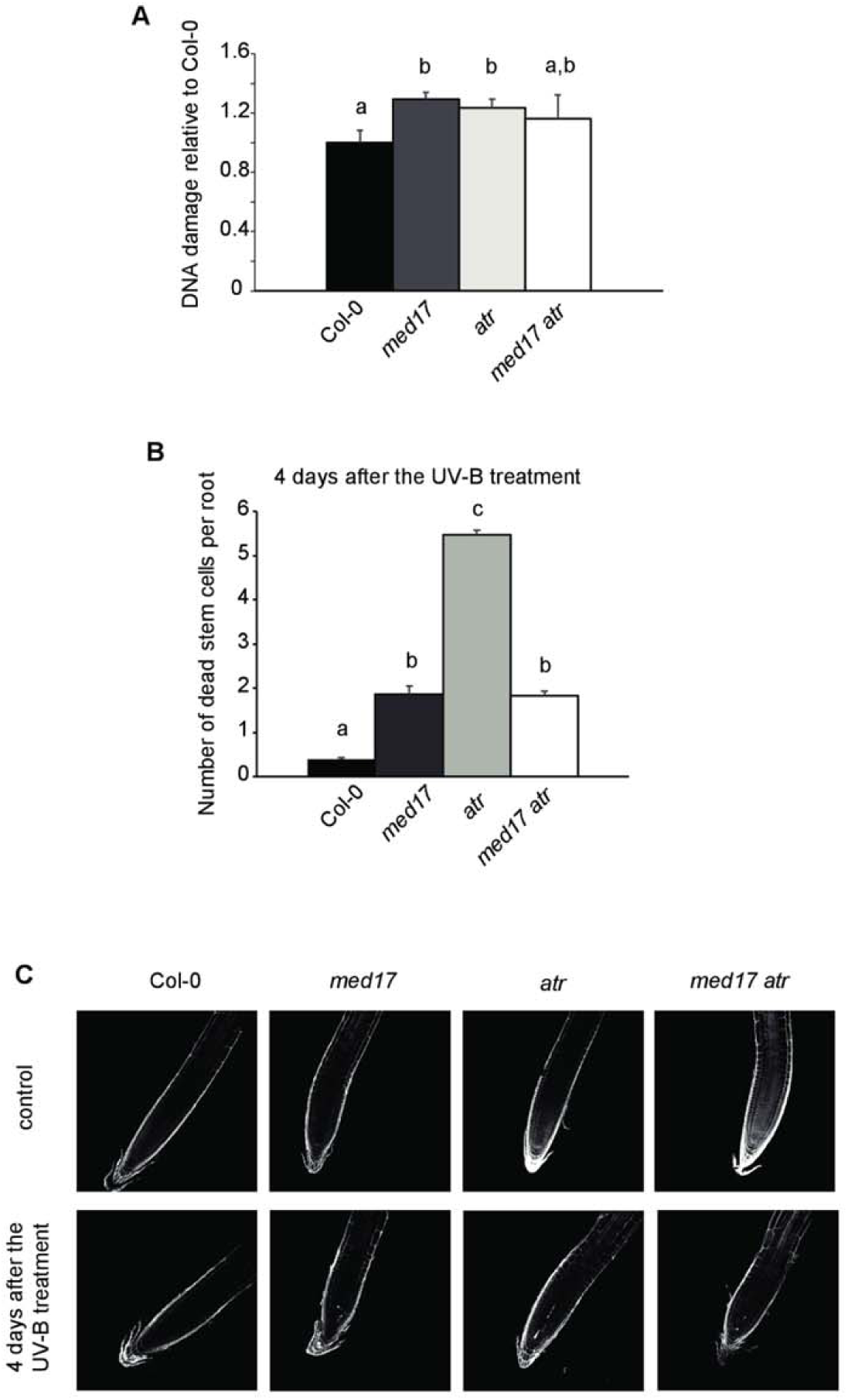
Characterization of DNA damage responses in double *med17 atr* mutant plants. A, Relative CPD levels in the DNA of WT Col-0, *med17, atr* and *med17 atr* plants immediately after a 4-h UV-B treatment under light conditions. Results represent averages ± S.E.M. of six independent biological replicates. B, Programmed cell death in meristematic root cells in Col-0, *med17*, *atr* and *med17 atr* plants 4 days after UV-B exposure. Results represent the average of at least 50 biological replicates ± S.E.M. Different letters indicate statistically significant differences applying analysis of variance test (*p* <0.05). C, Representative images of stem cells and adjacent daughter cells from WT Col-0, *med17, atr* and *med17 atr* seedlings that were scored for intense PI staining to count dead stem cells per root 4 days after a UV-B treatment or under control conditions.

The effect of UV-B was investigated on the meristematic zone of the primary roots. The size of the meristematic zone of the primary roots from *atr* seedlings was similar to that of WT primary roots under control conditions in the absence of UV-B, while the size of that in the double mutant was significantly smaller (Figure 7a). In UV-B treated roots, there was a significant decrease in the meristematic zone size in all lines; however, the decrease observed was significantly higher in *atr* mutants, while *med17 atr* plants showed a similar decrease as that in *med17* plants (Figure 7, A and B). This higher decrease in the meristem size in *atr* plants was a consequence of a higher decrease in the cortex cell number in the root meristem than that in *med17*, *med17 atr* and WT plants by UV-B, while all lines showed a similar increase in the cell area after the treatment (Figure 7, C-F). Similarly as for all other parameters analyzed, MED17 is required for the higher inhibition of cell proliferation in the meristematic zone of the primary roots observed in *atr* mutants under UV-B conditions. Therefore, *atr* phenotypes are suppressed in the absence of MED17. Interestingly, *med17* seedlings show increased *ATR* expression (Fig 6f); thus, the similar phenotypes observed in *med17* and *med17 atr* double mutants are independent of *ATR* levels.

**Figure 7.**
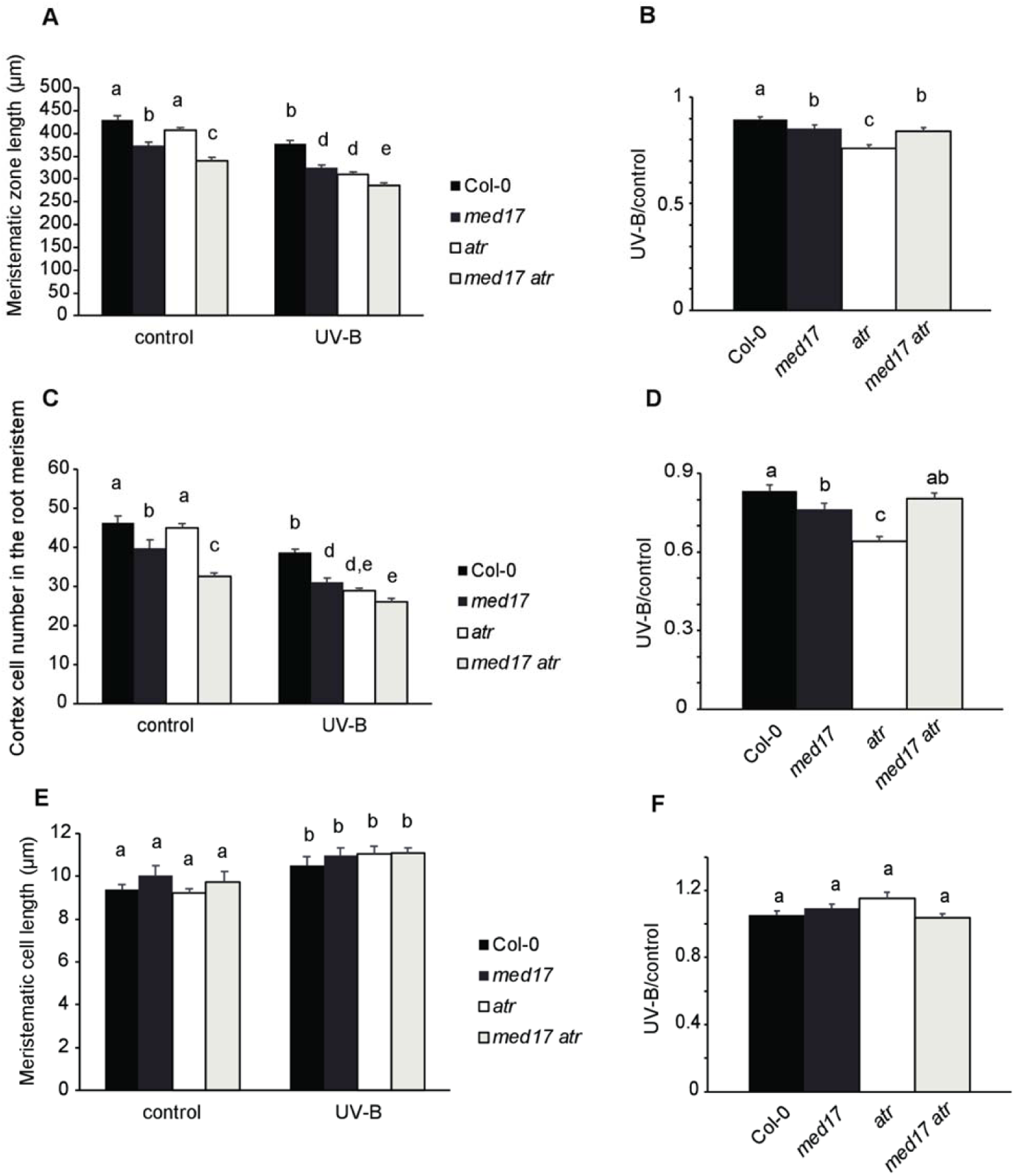
UV-B root meristematic zone of *med17 atr* is similarly affected by UV-B as *med17 s*eedlings but differently than *atr* mutants. A, Average of meristematic root zone length; C, cortex cell number; E, cortex cell length in the root meristem from WT Col-0, *med17*, *atr* and *med17 atr* seedlings after 4 days of a UV-B treatment or under control condition. B, D and F, Ratio between meristematic root zone length (B), cortex cell number (D), and cortex cell area values (F) measured after UV-B exposure vs those under control conditions are shown. Results represent the average ± S.E.M. Different letters indicate statistically significant differences applying analysis of variance test (P <0.05).

### MED17 deficiency is overcome by PDCD5 overexpression during DNA damage conditions after UV-B exposure

In addition, we investigated whether the role of MED17 in the DDR after UV-B exposure was affected in plants with altered expression of *PDCD5*. Previously, we showed that AtPDCD5 participates in the DNA damage response after UV-B exposure in Arabidopsis (Falcone Ferreyra et al., 2016). Plants that overexpressed AtPDCD5 accumulated lower levels of DNA damage after UV-B exposure and showed more PCD in root tips upon UV-B exposure (Falcone Ferreyra et al., 2016). Thus, we obtained transgenic plants that overexpressed PDCD5 in a *med17* mutant background by genetic crosses. When we analyzed DNA damage accumulation after UV-B in these plants, they showed lower amounts of CPDs than *med17* mutants, and even lower than WT plants, with CPD levels similar to those measured in *OE PDCD5* lines in a WT background (Figure 8, A). Moreover, when PCD was analyzed in the *OE PDCD5 med17* line one day after a UV-B treatment, these transgenic plants had a higher number of dead cells in the meristematic root zone than WT and *med17* roots, and similar to *OE PDCD5* plants in a WT background (Figure 8, B). Thus, after UV-B exposure, *PDCD5* overexpression counteracts the deficiency of *MED17* during the DDR.

**Figure 8.**
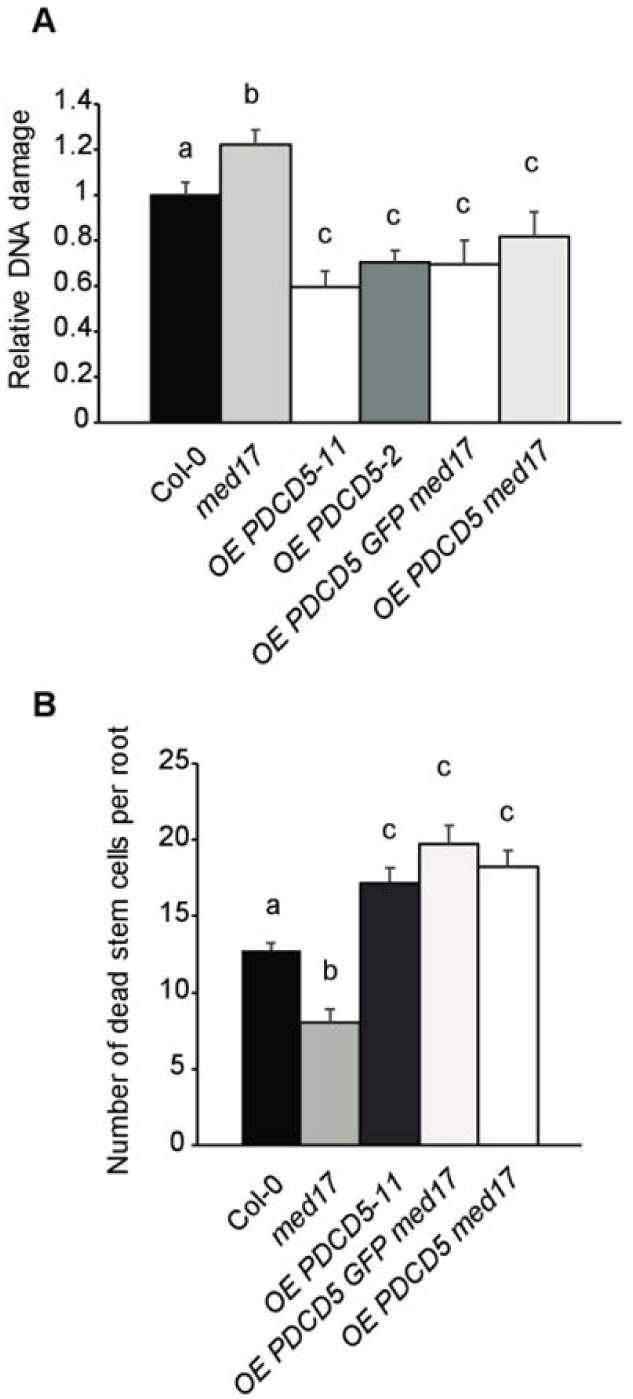
Characterization of DNA damage responses in *OE PDCD5 med17* plants. A, Relative CPD levels in the DNA of WT Col-0, *med17, OE PDCD5* and *OE PDCD5 med17* plants immediately after a 4-h UV-B treatment under light conditions. Results represent averages ± S.E.M. of six independent biological replicates. B, Programmed cell death in meristematic root cells of WT Col-0, *med17*, *OE PDCD5* and *OE PDCD5 med17* plants 1 day after UV-B exposure. Results represent the average of at least 50 biological replicates ± S.E.M. Different letters indicate statistically significant differences applying analysis of variance test (P <0.05).

When the expression of *PDCD5* was analyzed in *med17* and *OE MED17* seedlings, transcripts were significantly lower in *med17* than in WT after UV-B exposure, despite the opposite was observed under control conditions (Figure 4, L); while in *OE MED17* plants *PDCD5* expression was significantly higher than in WT plants, both under control conditions and after UV-B exposure (Figure 5, D). In this way, some of the phenotypes of *med17* and *OE MED17* plants after UV-B could be the result of altered *PDCD5* expression.

## Discussion

MED17 is a subunit of the head module of the Mediator complex, it has an important role interacting with several other Mediator subunits, being a key scaffold component of the whole complex (Guglielmi et al., 2004; Cevher et al., 2014; Maji et al., 2019). In Arabidopsis, MED17 is required for smRNA biogenesis and for the repression of heterochromatic loci, suggesting that this Mediator subunit, besides having a role in transcription, it would also participate in genome stability (Kim et al., 2011). In this work, we aimed to investigate the role of AtMED17 in UV-B responses. The analysis of the transcriptome profile of *med17* mutants compared to that of WT plants grown under white light conditions showed that of the 6822 genes that showed altered expression in *med17* mutants, 32% were also UV-B regulated in WT Col-0 plants reported in previous experiments (Tavridou et al., 2020). Of these 2184 genes, about 56% showed low expression in *med17* mutants and were up-regulated by UV-B in WT plants, suggesting that MED17 has a positive role in the regulation of UV-B responsive genes in Arabidopsis plants. In this group and as shown in Supplemental Figure S1, C, we found *RUP1* and *RUP2*, which encode two highly related WD40-repeat proteins that are negative regulators of the UVR8 photoreceptor in UV-B photomorphogenic responses (Gruber et al., 2010). *HY5* and *HYH,* which encode transcription factors that mediate UV-B responses of the UVR8-dependent pathway and activate the expression of *RUP1* and *RUP2,* also showed decreased expression in *med17* mutants. CRY1 and CRY3, which are blue light photoreceptors in the nuclei (CRY1) and in the chloroplasts and mitochondria (CRY3; Liu et al., 2011), were also down-regulated in *med17* plants. In particular, CRY3 belongs to the CRY-DASH clade of the photolyase/cryptochrome superfamily, and besides acting as a photoreceptor, it may also have single-strand DNA repair activity (Selby and Sancar, 2006; Pokorny et al., 2008). Interestingly, *UVR3*, which encodes an enzyme with 6–4 photolyase activity in Arabidopsis (Nakajima *et al*., 1998), increased in response to UV-B in WT, but not in *med17* plants, and was constitutively highly expressed in *OE MED17* plants. On the other hand, other genes that encode DNA recombination and repair, such as *RADA* and *RAD23 B-D* (Ishibashi et al., 2006; Lahari et al., 2018) showed up-regulation in *MED17* deficient plants, demonstrating that MED17 is required for proper expression of DNA repair enzymes.

In yeast, MED17 physically interacts with Rad2/XPG and participates in DNA repair after UV exposure, recruiting Rad2 to transcribed genes (Eyboulet et al., 2013). Moreover, in human cells, MED17 interacts with a DNA helicase XPB subunit of TFIIH, which is essential for both transcription and NER, and similarly as it was described in yeast, MED17 plays an important role switching between transcription and DNA repair (Kikuchi et al., 2015). After immunoprecipitation studies using transgenic plants expressing *MED17-GFP*, we demonstrated that in Arabidopsis, MED17 also directly or indirectly interacts with proteins that participate in DNA repair, such as the histone chaperones NAP 1; 3 and NRP 1 and 2 (Casati and Gomez, 2021) and a DNA repair ATPase-related protein and a replication factor C subunit 4 (Chen el al., 2018). Moreover, the results presented here show that *med17* mutants accumulate higher DNA damage than WT plants, while plants that overexpress *MED17* accumulate lower amounts of CDPs after UV-B exposure. In this way, and similarly as MED17 from yeast and humans, AtMED17 has a role in DNA repair. In yeast and humans, MED17 participates in transcription-coupled DNA repair (TCR) by NER DNA repair pathway, which removes DNA lesions that interfere with the progression of the RNA polymerase through transcribed genes (Eyboulet et al., 2013; Kikuchi et al., 2015; Hanawalt and Spivak, 2008). In Arabidopsis, *med17* plants are deficient not only in dark repair, which in plants is mostly achieved by NER, but also during light conditions, mostly accomplished by photolyases (Spampinato, 2017). Our results show that AtMED17 associates in a same complex with the transcription initiation factors TFIID subunit 9 and TFIIE subunit alpha, which may participate in TCR repair. Additionally, MED17, directly or indirectly interacting with the Replication Factor C subunit 4, could have a role in DNA repair by the NER system in Arabidopsis. AtMED17 also co-immunoprecipitated with histone chaperones and other chromatin associated factors, which may be required for proper dark DNA repair but also during photoreactivation. Thus, *med17* plants may be deficient in DNA repair due to decreased TCR repair and because MED17 may be necessary to interact with chromatin proteins during DNA repair. In addition, *med17* plants may accumulate more DNA damage after UV-B exposure because they express lower levels of some photolyases and other DNA repair proteins.

In addition, *med17* mutants showed a higher inhibition of cell proliferation in the root meristems after a UV-B treatment, and the meristems had less dead cells 1 day after, while they still presented dead cells after 4 days, in contrast to WT primary roots which showed more dead cells 1 day after the treatment but completely recovered after 4 days. These results suggest that MED17 may also participate in other aspects of the DDR besides DNA repair. A low accumulation of dead cells persisting after exposure under genotoxic conditions was also previously observed in *sog1* mutants (Johnson et al., 2018). SOG1 is a transcription factor that, in *Arabidopsis thaliana*, is a master regulator of genes that participate in the DNA damage response, including in the activation of programmed cell death in the meristematic cells in the roots. Johnson et al. (2018) demonstrated that, in contrast to WT plants, *sog1* mutants are defective in damage-induced programmed cell death and fail to undergo cell division in the primary roots meristems. A similar response was also observed in *med17* mutants in our experiments. Interestingly, *SOG1* transcript levels were significantly decreased in *med17* plants, suggesting that the PCD phenotype and the inhibition of cell proliferation in the primary root meristems after UV-B exposure in the *MED17* deficient plants can be due to decreased expression of this transcription factor.

Interestingly, other Mediator subunits have been shown to participate in PCD and root development, for example MED18, which is also a subunit of the Mediator head. *med18* mutants show a reduction in primary root growth, with an increase in lateral root formation and root hair development (Raya-Gonzalez et al., 2018). *med18* roots had altered cell division and elongation with an increased auxin response and transport at the root tip. Moreover, *med18* seedlings showed PCD in the root meristem in the absence of any genotoxic stress, which increased with age and/or exposition to DNA-damaging agents (Raya-Gonzalez et al., 2018). Despite MED17 and MED18 are both components of the Mediator head, they have different roles at least during DNA damage conditions. For example, while *med18* roots show high PDC in the meristematic primary root zone even in the absence of any genotoxic agent; *med17* roots show an opposite phenotype, with very low number of dead cells after UV-B exposure, and undetectable dead cells under control conditions in the absence of UV-B. Moreover, while meristematic cells in the primary roots of *med18* seedlings are bigger than those in WT roots (Raya-Gonzalez et al., 2018); cells in the meristematic zone of *med17* plants are similar to those in WT plants. *med17* root meristems are shorter than WT root meristems because they have less cells, and they show a higher decrease in the number of cells after UV-B exposure, suggesting that while MED18 may have an important role controlling cell size, MED17 may mostly control cell proliferation.

In Arabidopsis, both ATR and/or ATM regulate DNA damage responses after UV-B exposure (Furukawa et al., 2010). The double *med17 atr* mutant showed similar phenotypes as those of *med17* mutants, both under control conditions and after UV-B exposure; thus, MED17 is required for a proper activation of the DNA response mediated by ATR. It is possible that MED17 could act upstream ATR, for example interacting with DNA repair proteins during DNA damage recognition. The absence of proper damage recognition due to MED17 deficiency may therefore affect the activation of the DDR through ATR. However, we cannot rule out that MED17 may modulate ATR activity in other ways, for instance regulating the expression of proteins required for ATR activation of the DDR.

We previously characterized a UV-B inducible protein, AtPDCD5, which is similar to human PDCD5, a PCD-associated protein (Falcone Ferreyra et al., 2016; Xu et al., 2009). In humans, PDCD5 interacts with a histone acetyltransferase of the MYST family, TIP60, which are together recruited to chromatin in response to DNA damage, where they participate in different stages of repair (Murr et al., 2006; Xu et al., 2009). On the other hand, PDCD5 is also involved in the activation of PCD in the cytosol (Zhuge et al., 2011). *pdcd5* mutants accumulate higher levels of CPDs than WT plants after UV-B exposure but lower PCD in the primary root tip; these phenotypes are similar to those of *med17* mutants. On the contrary, plants overexpressing *AtPDCD5* were less sensitive to DNA damage and showed more dead cells in the root meristems after UV-B exposure (Falcone Ferreyra et al., 2016). Our results show that overexpression of *PDCD5* counteracts the deficiency in MED17 levels. As *PDCD5* levels are affected in plants with altered *MED17* expression, some of the phenotypes after UV-B exposure in *med17* and *OE MED17* plants could be, at least in part, the result of altered *PDCD5* expression.

In summary, our results demonstrate that MED17 regulates different plant responses to UV-B in Arabidopsis plants, in particular the DDR. According to the presented data, MED17 not only transcriptionally modulates the expression of genes of the DDR and the UV-B pathway, but it also physically interacts with transcription initiation factors and/or chromatin proteins that could facilitate DNA repair. Finally, we here show that MED17 is required for the *atr* mutant phenotypes, and that its deficiency is overcome by *PDCD5* overexpression. The interaction of MED17 with ATR and PDCD5 during the DDR may be regulating gene expression of proteins in the pathway, but it may also be during the early recognition of DNA damage through the binding with DNA repair proteins, which may be necessary for the activation of the pathway.

## Materials and methods

### Plant material, growth conditions and UV-B treatments

*A. thaliana* ecotype Col-0 was used for all experiments. *med17-1* (SALK_102813) mutants were provided by Dr. Xuemei Chen (University of California, Riverside, USA). *atr-2* (SALK_03284) and *atm-2* (SALK_006953) seeds were provided by Dr. Roman Ulm (University of Geneva, Switzerland). *OE PDCD5* transgenic plants (Falcone Ferreyra et al., 2016), *atr-2* or *atm-2* single mutants were crossed with *med17* mutants and the F2 population was screened by PCR using specific primers for *MED17*, *ATR, ATM* and *PDCD5* genes (Supplemental Table S4). For all experiments, F3 plants were used.

Arabidopsis plants were sown on soil, stratified for 3 days at 4 °C and they were then moved to a growth chamber. Plants were grown at 22 °C under a 16-h-light/8-h- dark photoperiod (100 μE m^−2^s^−1^). For root and hypocotyl studies, plants were germinated and grown on petri plates containing Murashige and Skoog salt (MS)-agar (0.7 % w/v) medium for 5 days.

For all UV-B treatments except for flowering time assays, plants were irradiated with UV-B lamps using fixtures mounted 30 cm above the plants (2 W m^−2^ UV-B and 0.6 W m^−2^ UV-A, Bio-Rad ChemiDoc™XRS UV-B lamps, catalogue 1708097). The lamps have emission spectra from 290 to 310 nm, with a maximum emission peak at 302 nm. The bulbs were covered using cellulose acetate filters (100 mm extra-clear cellulose acetate plastic, Tap Plastics, Mountain View, CA); the cellulose acetate filters absorb wavelengths lower than 290 nm without removing UV-B from longer wavelengths. As a control treatment without UV-B, plants were exposed for the same period of time under lamps covered with a polyester plastic that absorbs UV-B at wavelengths lower than 320 nm. For root and hypocotyl elongation assays, seedlings were irradiated for 1 h. For DNA damage analysis, 4-week-old plants were irradiated with UV-B for 4 h, and leaves were collected immediately and after 2 h of recovery in the absence of UV-B, either under light or dark conditions. UV radiation was measured using a UV-B/UV-A radiometer (UV203 AB radiometer; Macam Photometrics).

For flowering time analysis, white light was supplemented with 2 Wm^-2^ of UV-B (311 nm; Phillips narrowband TL/01 lamps) during 1 h every day starting from day 9 after transferring to the growth chamber until flowering, the zeitgeber time of UV-B treatments was 4 h in long day conditions.

For seedling lethality analysis, seeds were sown in agar plates and stratified for 3 days. Then, they were irradiated with white light (100 μE m^−2^ s^−1)^ for 1 h and, after that, they were kept 24 h in the dark. Next, plates were treated with UV-B for 1 h, transferred to darkness for 48 h and then they were finally allowed to grow in the growth chamber for 15 days under normal light conditions (UV-B irradiated plants). Alternatively, a different group of seedlings were grown under the same conditions but they were not UV-B irradiated (darkness treated plants). Additionally, seedlings were stratified for 3 days and they there were grown under normal growth conditions for 18 days (light grown plants).

UV radiation was measured using a UV-B/UV-A radiometer (UV203 AB radiometer; Macam Photometrics).

### DNA damage analysis

12 DAS leaf samples from plants treated with UV-B or kept under control conditions were collected immediately or 2 hours after the end of the treatment and immersed in liquid nitrogen. CPD accumulation in the DNA purified from the collected samples was analyzed as described previously (Lario et al., 2013). UV-B treatments were performed both under light and dark conditions; plants irradiated under dark conditions were allowed to recover for 2 h under light or dark conditions. 0.1 g were collected and extracted DNA was dot blotted onto a nylon membrane (Perkin-Elmer Life Sciences). The blot was incubated with monoclonal antibodies specific to CPDs (TDM-2) from Cosmo BioCo (1:2,000 in TBS). Quantification was achieved by densitometry of the dot blot using Image-Quant software version 5.2.

### Root meristem analysis and programmed cell death after UV-B exposure

Seedlings were grown for 5 days in vertically oriented Murashige and Skoog plates, and were then irradiated with UV-B light or kept without UV-B. UV-B-irradiated and control seedlings were then incubated for 24 hr or 96 hr in the growth chamber, and then PCD was analyzed as described by Furukawa et al. (2010). Root tips were stained using a modified pseudo-Schiff propidium iodide staining protocol and visualized by confocal laser scanning microscopy (Nikon C1) under water with a 40× objective. The excitation wavelength for propidium iodide-stained samples was 488 nm, and emission was collected at 520 to 720 nm. Dead (intensely Propidium iodide (PI)-staining) cells in the vicinity of the quiescent centre were counted and scored as dead cells per root.

### Generation of Arabidopsis transgenic plants

cDNA was obtained from leaf tissues of WT plants grown under continuous light growth. *MED17* cDNA without its stop codon was amplified using specific primers including *KpnI* and *XhoI* restriction sites (Table S1). PCR was done using Pfu (Invitrogen) polymerase under the following conditions: 94°C for 5 min; 40 cycles of 94°C for 30 sec, 55°C for 20 sec and 72°C for 30 sec; and finally, one cycle at 70°C for 2 min. PCR product was purified from the gel, cloned in a pBluescript vector and sequenced. The construct was then transformed into *E. coli* DH5α and then the plasmid was purified and digested with *KpnI* and *XhoI*. The digestion fragment corresponding to *MED17* was subcloned into the pCardo plasmid, and the construct expressing *MED17* under the *35S* promoter was transformed into *Agrobacterium tumefaciens* strain GV3101. Col 0 was transformed using the floral dip method (Clough and Bent,1998). In addition, the pBluescript vector with *MED17* cDNA was digested using *Kpn* and *SalI* and the fragment was cloned into pCS052_GFP_pCHF3 (a modified version of pCHF3; with the GFP coding sequence without the start codon inserted into *SalI-PstI* sites). The resulting construct, pCHF3:*MED17-GFP,* was transformed into *E. coli* DH5α and purified. The construct was transformed into *Agrobacterium tumefaciens* GV3101 and Col 0 and *med17* plants were transformed using the floral dip method. Transformed seed (T1) were identified by selection on solid MS medium containing kanamycin (30 mg L^-1^, pCHF3) or BASTA (3 mg mL^-1^, pCardo, and finally plants were transferred to soil. The presence of *Pro35S:MED17 (OE MED17)* and *Pro35S:MED17-GFP (OE MED17-GFP)* transgenes in T2 plants was screened by PCR using genomic DNA (Table S1).

### Flowering time analysis

Flowering time was determined by counting the number of rosette leaves or the number of days until the first flower opens, similar to previous reports (Dotto et al., 2018). Flowering time was counted as the number of rosette leaves at the moment of flowering or the number of days until the first flower opens.

### Seed analysis

Seeds were phenotypically analyzed using a Nikon SMZ-10 microscope. For silique analyzes, 56 siliques from each genotype were analyzed, and the number of aborted seeds per silique were counted. After that, seeds were stained using PI stain and they were then observed using confocal laser scanning microscopy, using a Confocal Nikon C1 microscope.

### RNA-seq experiments

For the RNA-seq experiments, WT and *med17* seeds were sown on MS medium with 0.8 % (w/v) agar, kept at 4°C for three days and then grown for 10 days at 23°C under long day (16 h light, 8 h dark, 100 μE m^−2^ s^−1^) of white fluorescent light. Three independent biological replicates for each genotype were harvested 2 h before the start of the night period and the seedlings were immediately frozen in liquid nitrogen. Total RNA was prepared using a Plant Total RNA Mini Kit (YRP50).

FASTQC v0.11.5 was used for quality control of the FASTQ sequence files (Andrews, 2010). Illumina 150-bp paired-end reads were mapped to the *A. thaliana* reference genome assembly (assembly version TAIR10) with HISAT2 (Kim et al., 2015) and raw read counts per gene were then estimated with htseq-count (Anders et al., 2015) and normalized according to trimmed mean of M-values (TMM) (Robinson and Oshlack, 2010). Over 22 million reads were obtained for each sample, with an overall alignment rate of 92%. Genes with more than five reads per million in only two or fewer samples were eliminated from the analysis. Differential expression analysis of the remaining genes was carried out with the R Package EdgeR (Robinson et al., 2010) using a quasi-likelihood negative binomial generalized log-linear model (EdgeR function glmQLFit) (Lun et al., 2016). Genes with FDR<0. 05 were selected. A complete list of identified transcripts is in Supplemental Table S1. Venn diagrams were generated with VennDiagram R package (Chen and Boutros, 2011), and the Heatmap was generated with Complex Heatmap R package (Gu et al., 2016). GO analysis was performed using DAVID bioinformatics resources (Huang et al., 2009).

RNA seq data from Arabidopsis Col-0 plants UV-B irradiated was obtained from Tavridou et al. (2020). FASTQ files were obtained from Gene Expression Omnibus (GEO) repository, and processed as med17 files. Statistical analysis of the overlapping differentially expressed genes was done using the R Package GeneOverlap.

### qRT-PCR analysis

Analysis was done as described in Maulion et al. (2019). Briefly, Total RNA was isolated using the TRIzol reagent (Invitrogen). 0.5 to 1.0 mg of total RNA was reverse transcribed using SuperScript II reverse transcriptase (Invitrogen) and oligo (dT) as a primer. The resultant cDNA was used as for quantitative PCR amplification in a StepOne™ System apparatus (ThermoFisher Scientific), using SYBRGreen I (Invitrogen) as a fluorescent reporter and Platinum taq polymerase (Invitrogen). Transcript levels were normalized to those of the *A*. *thaliana* calcium-dependent protein kinase3 (Supplemental Table S4) and to values in Col-0 plants grown under control conditions in the absence of UV-B.

### Coimmunoprecipitation studies and MS analysis

For coimmunoprecipitation analyses, 3 g of Arabidopsis leaves were homogenized in a buffer containing 0.4M sacarose, 10mM Tris-HCl, pH 8.0, 10mM MgCl_2_ and 1mM phenylmethylsulfonyl fluoride (PMSF). The extract was filtered through Miracloth and next was centrifugated for 20 min at 4500xg. The pellet was resuspended in buffer 2 (0.25M sacarose, 10mM Tris-HCl, pH 8.0, 10mM MgCl_2_, 0.15% (v/v) Triton X-100, 5mM 2 mercaptoethanol and 0.1 mM PMSF). After 5 min of incubation in ice, the extract was centrifuged at 5000xg for 10 min. The pellet was resuspended in buffer 3 containing 0.44 M sacarose, 25mM Tris-HCl, pH 7.6, 10mM MgCl_2_, 0.5% (v/v) Triton X-100 and 10mM 2-mercaptoethanol. The supernatant was discarded, and the pellet was resuspended in buffer 4 (0.44 M sacarose, 50mM Tris-HCl, pH 7, 5mM MgCl_2_, 20% (v/v) glycerol and 10mM 2-mercaptoethanol) and centrifuged for 10 min at 12500xg. Finally, the pellet was resuspended in 200 µl of lysis buffer (10mM Tris-HCl, pH 7.5, 50mM NaCl, 0.1% (v/v) Triton X-100, 10% (v/v) glycerol and 1mM PMSF) and sonicated. Then, the extract was centrifugated at 17500xg for 15 min. After centrifugation, 1 mL of crude extract (0.75 mg of total protein) was incubated with 6 µL (3 mg) of affinity-purified rabbit polyclonal antibody raised against GFP for 3 h at 4°C with gentle agitation. After this, 20 µL of protein A agarose was added, and the samples were incubated at 4°C with gentle agitation for 1 h. The agarose beads were pelleted by centrifugation and washed four times with 200 µL of lysis buffer (100 mM Tris-HCl, pH 7.5, 1 mM EDTA, 150 mM NaCl, and 1% (v/v) Triton X-100) for 5 min at 4°C and once with LNDET buffer (250 mM LiCl, 1% Nonidet P-40, 1% [w/v] deoxycholic acid, 1 mM EDTA, and 10mM Tris-HCl, pH 8.0) for 5 min at 4°C. Proteins were eluted by incubation at 95°C for 5 min in 50 μL of SDS sample buffer. Samples were loaded on 10% SDS-PAGE gels and run at 100 V for 15 min to allow proteins to migrate less than 1 cm into the resolving gel. Gels were stained with colloidal Coomassie Brilliant Blue stain, and the immunoprecipitated protein-loaded lane was cut into one rectangular slice of less than 1 cm of height.

The gel slices were subjected to in-gel digestion (Link and LaBaer, 2009; http://cshprotocols.cshlp.org/content/2009/2/pdb.prot5110.abstract) with trypsin (porcine, side chain protected; Promega). Briefly, specific excised samples were washed once with 50% (v/v) acetonitrile in 50 mM NH_4_HCO_3_ and then dehydrated with pure acetonitrile. The gel samples were next reduced with dithiothreitol (DTT; 10 mM in 25 mM NH_4_HCO_3_, 65°C for 30 min) and alkylated with iodoacetamide (55 mM in 25 mM NH_4_HCO_3_, room temperature for 30 min). Then, the gel pieces were incubated with acetonitrile, and rehydrated in 50 uL of digestion buffer (12 ng mL trypsin in 25 mM NH_4_HCO_3_) After overnight digestion at 37°C, peptides were extracted once with a solution containing 66% (v/v) acetonitrile and 5% (v/v) formic acid. The supernatants were concentrated to 5 µL by centrifugation under vacuum. The digests were analyzed by capillary HPLC-MS/MS.

### HPLC-MS/MS

The peptide mixtures were analyzed in data-dependent mode on a Q-Exactive HF mass spectrometer coupled to an Ultimate 3000 nanoHPLC. A volume of 4 μl of peptide samples was loaded by the LC system. Peptides were desalted online on a reverse-phase C18 cardtridge using buffer A (0.1% (v/v) formic acid) as running buffer, and then resolved on a 15-cm long PepMap nanocolumn (EASY-Spray ES801, Thermo) at a flow rate of 0.3 μl/min. Peptide elution was achieved with a gradient of buffer B (100% acetonitrile containing 0.1% formic acid). Total run time was 150 min and programmed as follows: 15 min column equilibration in 96% buffer A, 4% buffer B, followed by a 100 min gradient from 4% buffer B to 35%. Then, a steeper gradient from 35% buffer B to 90% was carried out in 25 min. 90% buffer B was maintained for 5 min and finally, the system was allowed to reach initial conditions in 5 min.

For mass spectrometric analysis in the Q Exactive HF mass spectrometer, the following tune method was used: full scan spectral range from *m/z* 375 to 1600, automatic gain control (AGC) target value set at 3 × 10^6^, and a mass resolving power of 120,000 for full spectra. MS/MS were analyzed in data-dependent mode with a resolution of 30,000 and an AGC target of 5 x 10^5^. Up to 20 precursors were selected for dissociation in the high-energy collisional dissociation chamber using a normalized collision energy of 27. Ion selection was performed applying a dynamic exclusion window of 15 sec.

For protein identification, all raw LC−MS/MS data were analyzed by MaxQuant v1.6.17.0 using the Andromeda Search engine and searched against the *A. thaliana* database downloaded from Uniprot (August 2020 release with 39,346 protein sequences). Parameters for MS/MS spectra assignment were as follows: full-trypsin specificity, maximum of two missed cleavages, instrument default parameters set for Orbitrap, carbamidomethylated cysteine as a fixed modification, and oxidized methionine and N-acetylation of protein termini as variable modifications. False discovery rate at both peptide and protein levels was set to 1%. Data filtering, processing and interpretation were performed in Perseus v1.6.14.0.

### Quantification of UV absorbing compounds

One-half gram of fresh leaf tissue was frozen in liquid nitrogen and ground to a powder with a mortar and pestle. The powder was extracted for 8 hr with 3 mL of acidic methanol (1% HCl in methanol), and then by a second extraction with 6 mL of chloroform and 3 mL of distilled water. The extracts were vortexed and then centrifuged 2 min at 3,000×*g*. UV-B absorbing compounds were quantified by absorbance at 330 nm.

### Statistical analysis

Statistical analysis was done using analysis of variance models (Tukey test) or alternatively Student’s *t* test (Welch’s *t* tests), using untransformed data.

## Accession numbers

Sequence data from this article can be found in the The Arabidopsis Information Resource under accession number At5G20170.

## Acknowledgments

M.L.F.F., P.Ce. and P.Ca. are members of the Researcher Career of the Consejo Nacional de Investigaciones Científicas y Técnicas (CONICET). M.L.F.F. and P.Ca. are Professors at UNR. M.S.G is a doctoral fellow from CONICET, and M.S. is a doctoral fellow from FONCYT. We thank María José Maymó (CEFOBI) for care in cultivating Arabidopsis plants and Mariana Giro for help in confocal microscope imaging.

## Supplemental data

**Supplemental Figure S1** Venn diagrams of comparisons between transcripts with altered expression in *med17* mutants and UV-B-responsive genes in Arabidopsis plants.

**Supplemental Figure S2** Analysis of *med17* and *OE med17* plants after UV-B exposure.

**Supplemental Figure S3** Representative pictures of individual WT Col-0, *med17* and *OE MED17* plants.

**Supplemental Figure S4** UV-B effect on cell proliferation in the root meristematic zone of WT Col-0, *med17* and *OE MED17* seedlings 4 days after UV-B exposure.

**Supplemental Figure S5** UV-B effect on the root meristematic zone of WT Col-0 and *med17* 1 day after UV-B exposure.

**Supplemental Figure S6** Characterization of double *med17 atr* mutant plants.

**Supplemental Figure S7** Confocal microscopy of PI stained seeds from WT Col-0, *med17*, *atr* and *med17 atr* plants.

**Supplemental Figure S8** Programmed cell death in meristematic root cells in WT Col-0, *med17*, *atr* and *med17 atr* plants 1 day after UV-B exposure.

**Supplemental Table S1** List of transcripts expressed in *med17* mutants and comparison with expression levels in WT Col-0 plants.

**Supplemental Table S2** Cluster and GO analysis of transcripts with differential expression both in *med17* mutants compared to WT plants and after UV-B exposure in WT Col-0 plants.

**Supplemental Table S3** List of potential MED17 interaction proteins enriched in the MED17-GFP IP experiments.

**Supplemental Table S4** Primers used in for the experiments described in this work.

## Notes

**Funding** This research was supported by Argentina FONCyT grants PICT 2016-141 and 2018-798

